# Local computational methods to improve the interpretability and analysis of cryo-EM maps

**DOI:** 10.1101/2020.05.11.088013

**Authors:** S. Kaur, J. Gomez-Blanco, A. Khalifa, S. Adinarayanan, R. Sanchez-Garcia, D. Wrapp, J. S. McLellan, K. H. Bui, J. Vargas

## Abstract

Cryo-electron microscopy (cryo-EM) maps usually show heterogeneous distributions of B-factors and electron density occupancies and are typically B-factor sharpened to improve their contrast and interpretability at high-resolutions. However, ‘over-sharpening’ due to the application of a single global B-factor can distort processed maps causing connected densities to appear broken and disconnected. This issue limits the interpretability of cryo-EM maps, i.e. *ab initio* modelling. In this work, we propose 1) approaches to enhance high-resolution features of cryo-EM maps, while preventing map distortions and 2) methods to obtain local B-factors and electron density occupancy maps. These algorithms have as common link the use of the spiral phase transformation and are called LocSpiral, LocBSharpen, LocBFactor and LocOccupancy. Our results, which include improved maps of recent SARS-CoV-2 structures, show that our methods can improve the interpretability and analysis of obtained reconstructions.

## Introduction

Cryo-electron microscopy (cryo-EM) has become a mainstream technique for structure determination of macromolecular complexes at close-to-atomic resolution and ultimately for building an atomic model (Ge, Scholl et al. 2020, Wandzik, Kouba et al. 2020). With its unique ability to reconstruct multiple conformations and compositions of the macromolecular complexes, cryo-EM allows the understanding of the structural and assembly dynamics of macromolecular complexes in their native conditions (Davis, Tan et al. 2016, Plaschka, Lin et al. 2017, Razi, Davis et al. 2019). However, the presence of heterogeneity in cryo-EM maps leads to a high variability in resolution within different regions of the same map. This leads to challenges and errors in the process of building an atomic model from a cryo-EM reconstruction. Additionally, current reconstructions from cryo-EM do not provide essential information to build accurate *ab initio* atomic models as atomic Debye-Waller factors (B-factors) or atomic occupancies, while their counterparts from X-ray crystallography do by analyzing the attenuation of scattered intensity represented at Bragg peaks.

Cryo-EM structures exhibit loss of contrast at high-resolution coming from many different sources, including molecular motions, heterogeneity and/or signal damping by the transfer function of the electron microscope (CTF). Interpretation of high-resolution features in cryo-EM maps is essential to understanding the biological functions of macromolecules. Thus, approaches to compensate for this contrast loss and improve map visibility at high-resolution are crucial. This process is usually referred to as “sharpening” and is typically performed by imposing a uniform B-factor to the cryo-EM map that boosts the map signal amplitudes within a defined resolution range. When the map is sharpened with increasing positive B-factors the clarity and map details initially improve, but eventually the map becomes worse as the connectivity is lost, and the map densities appear broken and noisy. In the global sharpening approach (Rosenthal and Henderson 2003, Fernandez, Luque et al. 2008, Scheres 2015), the B-factor is automatically computed by determining the line that best fits the decay of the spherically averaged noise-weighted amplitude structure factors, within a resolution range given by [15-10Å, *R*_max_], with *R*_max_ the maximum resolution in the map given by the Fourier Shell Correlation (FSC). More recently, the AutoSharpen method within Phenix (Terwilliger, Sobolev et al. 2018) calculates a single B-factor that maximizes both map connectivity and details of the resulting sharpened map. AutoSharpen automatically chooses the B-factor that leads to the highest level of detail in the map, while maintaining connectivity. This combination is optimized by maximizing the surface area of the contours in the sharpened map.

The approaches presented above are global, so the same signal amplitude scaling is applied to map regions that may exhibit very different signal to noise ratios (SNRs) at medium/high-resolutions. Thus, cryo-EM maps showing inhomogeneous SNRs (and resolutions) can result into sharpened maps that show both over-sharpened and under-sharpened regions. The former may be strongly affected by noise and broken densities, while the latter may present reduced contrast at high-resolutions. Both cases make it difficult or even impossible to interpret the biological relevance of these regions or even the whole map (Murshudov 2016). Thus, local sharpening methods have been proposed to overcome these limitations (Erney Ramirez-Aportela 2017, Jakobi, Wilmanns et al. 2017). LocScale approach (Jakobi, Wilmanns et al. 2017) compares radial averages of structure factor amplitudes inside moving windows between the experimental and the atomic density maps. After, the method modifies locally the map amplitudes of the experimental map in Fourier space to rescale them accordingly to those of the atomic map. This approach requires as input a complete atomic model (without major gaps) fitted to the cryo-EM map to sharpen, which is not always available. In addition, the size of the moving window should be provided and depending on the quality of the map to be sharpened, this process may lead to over-fitting. More recently, the LocalDeblur method (Erney Ramirez-Aportela 2017) proposed an approach for map local sharpening using as input an estimation of the local resolution. The method assumes that the map local density values have been obtained by the convolution between a local isotropic low-pass filter and the actual map. This local low-pass filter is assumed Gaussian shaped so that the frequency cutoff is given by the local resolution estimation.

In X-ray crystallography, the B-factor (also called temperature value or Debye-Waller factor) describes the degree to which the electron density is spread out, indicating the true static or dynamic mobility of an atom and/or the positions where errors may exist in the model building. The B-factor is given by 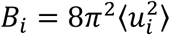, where 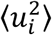 is the mean square displacement for atom *i*. These atomic B-factors can be experimentally measured in X-ray crystallography, introduced as an amendment factor of the structure factor calculations since the scattering effect of X-ray is reduced on the oscillating atoms compared to the atoms at rest (Sherwood, Cooper et al. 2011). B-factors can be further refined by model building packages, i.e. Phenix (Liebschner, Afonine et al. 2019) or Refmac (Winn, Murshudov et al. 2003) to improve the quality and accuracy of atomics models. Although B-factors are essential to ‘sharpen’ cryo-EM maps at high-resolution, they also provide key information to analyze cryo-EM reconstructions. Effective B-factors are used to model the combined effects of issues such as molecular drifting due to charging effects, macromolecular flexibility or possible errors in the reconstruction workflow that lead to a signal falloff (Rosenthal and Henderson 2003, Liao and Frank 2010, Penczek 2010). However, cryo-EM maps are usually analyzed with a single B-factor, even though maps may largely differ in different regions. Thus, methods to determine local B-factors are much needed to accurately analyze cryo-EM maps and improve the quality of fitted atomic models. Another local parameter usually provided by X-ray crystallography in contrast with cryo-EM are atomic occupancies (or Q-values). The occupancy estimates the presence of an atom at its mean position and it ranges between 0.0 to 1.0. Note that these parameters can be also refined by model building packages if the electron density map is of sufficient resolution. To our knowledge, currently there is not any available method to estimate local occupancies from cryo-EM maps, even though this information (in addition to local B-factors) is essential to building accurate atomic models. For example, in (Afonine, Klaholz et al. 2018) authors found that 31% of all models examined in this analysis possess unrealistic occupancies or/and B-factor values, such as all being set to zero or other unlikely values. They also reported that 40% of models analyzed show cross-correlations between cryo-EM maps and respective models below 0.5, and they indicated as a possible hypothesis an incomplete optimization of the model parameters (coordinates, occupancies and B-factors).

In this work, we propose semi-automated methods to enhance high-resolution map features to improve their visibility and interpretability. More importantly, these approaches do not require input parameters as fitted atomic models or local resolution maps, which reduces the possibility of overfitting. In particular, our proposed local map enhancement approach (LocSpiral) is robust to maps affected by inhomogeneous local resolutions/SNRs, thus the method strongly improves the interpretability of these maps. Secondly, we also propose approaches to determine local B-factors and density occupancy maps to improve the analysis of cryo-EM reconstructions. The link between the different proposed approaches is the use of the spiral phase transform to estimate a modulation or amplitude map of the cryo-EM reconstruction at different resolutions.

## Results

We tested our proposed methods with four different samples ranging from near-atomic single-particle reconstructions (∼1.54 Å) to maps with more modest resolutions (∼6.5 Å). In all cases, we compared our results with the ones provided by the Relion postprocessing approach (Fernandez, Luque et al. 2008, Kimanius, Forsberg et al. 2016).

### Polycystin-2 (PC2) TRP channel

First, we analysed a single-particle reconstruction of the polycystin-2 (PC2) TRP channel (EMDataBank: EMD-10418) (Wang, Corey et al. 2020). In this case, we focussed on showing the capacity of our LocSpiral approach, though, for the sake of consistency, we also show results of our obtained B-factor and occupancy maps. The original publication reports a resolution of 2.96 Å with a final B-factor to be used for sharpening of −84.56 Å^2^ (slope of Guinier plot fitting equal to −21.14 Å^2^). In Figure 1A, we show maps at different orientations and with high threshold values obtained by LocSpiral and by the postprocessing method of Relion 3 (Fernandez, Luque et al. 2008, Kimanius, Forsberg et al. 2016). The map densities are similar in the core of the protein, as can be seen from the red square in the figure where we show a zoomed view of the protein inner core or from the image with both maps superimposed. However, the map densities are quite different at the outer regions, where the Relion map shows thin and broken densities. In addition, we show comparisons of fitted densities with the corresponding atomic model (PDB ID: 6t9n) of two α-helices and one loop. The asterisks label results obtained by LocSpiral. The residues marked with a red arrow were used to adjust the threshold values between maps. These comparisons show that the map obtained by LocSpiral shows fewer fragmented and broken densities and a better coverage of the atomic model, helping in the interpretation of the maps and in the process of building accurate atomic models.

**Figure 1.**
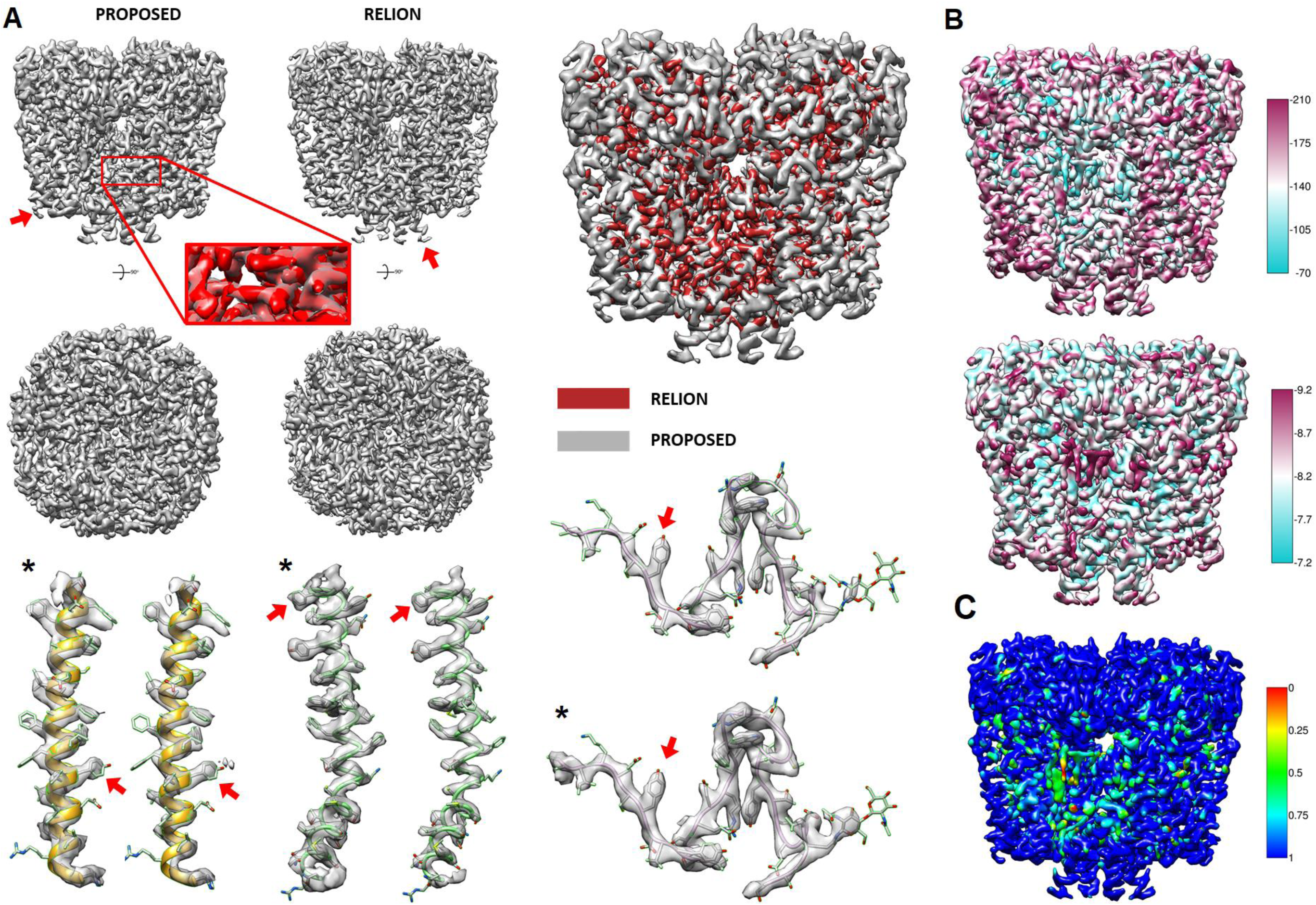
A) Top: Maps obtained by our local enhancement approach and by Relion postprocessing method at different orientations. The threshold values are adjusted to provide similar densities in the core inner part of the protein. The red square in the figure shows a zoomed view of the protein inner core where both maps (proposed and Relion) are superimposed. Relion map appears in red color, while our proposed map is in gray. Down: Fitted map densities (Proposed and Relion) with corresponding atomic model (PDB ID: 6t9n) of two α-helices and one loop. The asterisks mark our proposed approach. The residues marked with a red arrow were used to adjust the threshold values between maps. B) Obtained local B-factor map (upper map) and the A map that corresponds to the local values of the logarithm of structure factor amplitudes at 15 Å (y-intercept of the fit at 15 Å). C) Obtained local occupancy map.

We also compared the performance of LocSpiral with other methods, including LocalDeblur, our proposed local B-factor correction method (LocBSharpen) and the global B-factor correction approach as implemented in Relion. To compare the different results, we used metrics proposed in (Afonine, Klaholz et al. 2018). The results are shown in Figure S1. In this case, we used a relatively high threshold value to visually compare the different maps. From Figure S1, we can see that the map obtained by our proposed method shows good connectivity and is less affected by broken or missed densities. EMRINGER (Barad, Echols et al. 2015) and cross-correlation scores (obtained using PDB 6t9n as reference) show approximately similar results for all cases, though, the highest scores are provided by LocSpiral and LocBSharpen approaches. For the sake of comparison, we also provide FSC curves calculated by comparing the different maps with the reference atomic model (PDB 6t9n). In this case, the best results at high resolutions are provided by LocalDeblur and by our proposed LocSpiral approach.

In addition, we provide the results of our local B-factor and occupancy map calculations. In Figure 1B (upper map), we show the obtained local B-factor map to be used for sharpening (slope of the local Guinier plot multiplied by a factor 4) and the A map (local values of the logarithm of structure factors amplitudes at 15 Å). The resolution range used to estimate these maps was between 15 Å to the FSC resolution (2.96 Å). The average value of the local B-factor map gives a value of −89.45 Å^2^, which is in close agreement with the value provided by Relion (−84.56 Å^2^). The A map provides the fitted local amplitudes at 15 Å, showing the local “amount” of signal at this resolution. As expected, Figure 1B shows that the inner parts of the protein show lower B-factors than the outer regions. In Figure 1C, we show the obtained local occupancy map. Interesting, both the occupancy and A maps show low values in the regions occupied by detergent densities, lipid densities and cholesterol densities (please see Figure 2 in (Wang, Corey et al. 2020)), indicating the presence of compositional variability in these regions and low signal at 15 Å.

**Figure 2.**
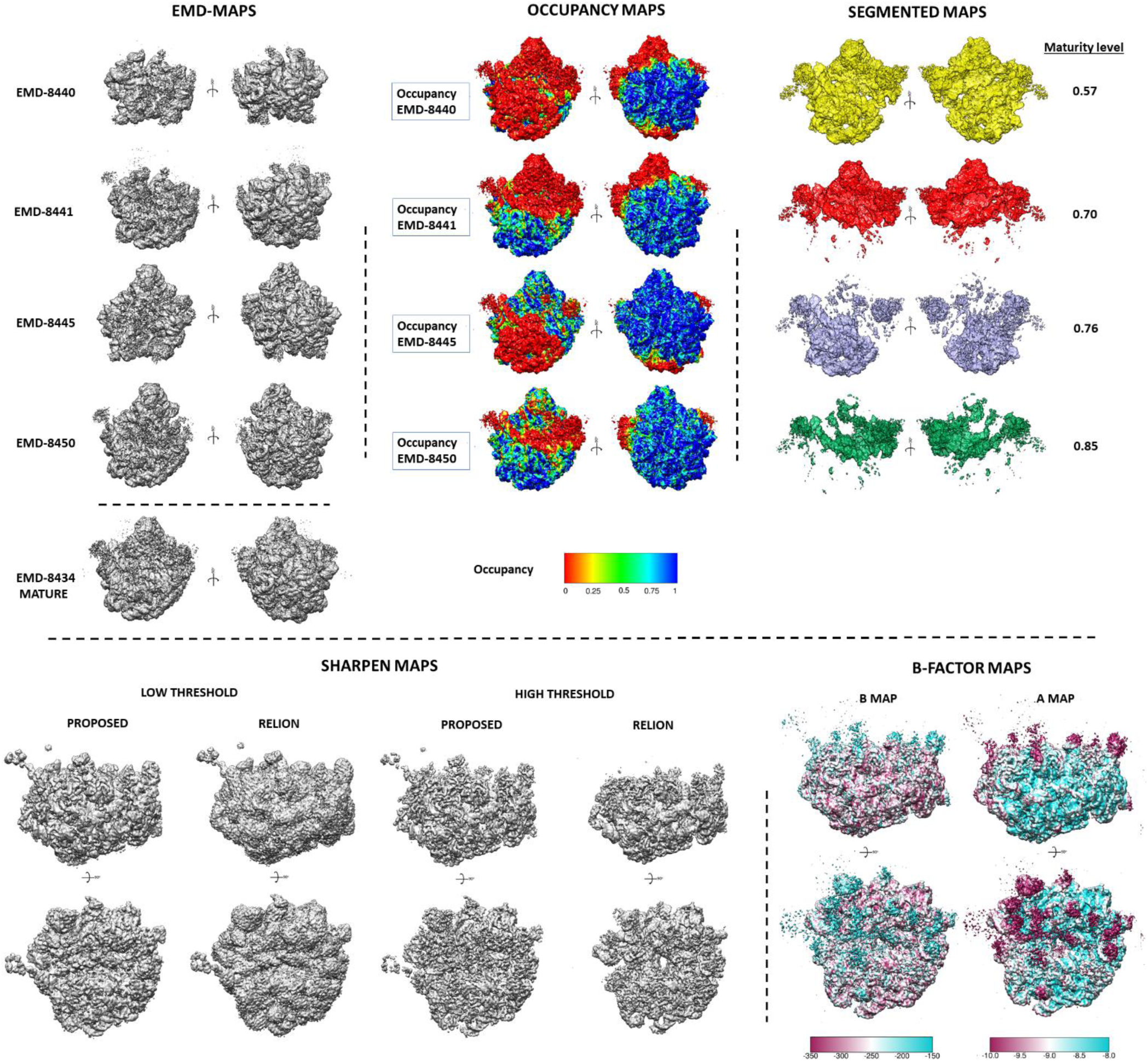
First row first column: different maps at different orientations as deposited in EMDB; Second column: obtained occupancy maps, where the mature 50S ribosome (EMDB-8434) is coloured with corresponding occupancy maps; Third column: Segmented maps showing the densities that are missing in the different immature maps when compared to the mature 50S reconstruction, and obtained maturity levels; Second row first column: Obtained improved maps of EMD-8441 by our proposed local enhancement approach and Relion. These maps are shown at low and high thresholds; Second row second column: Obtained B-maps (local B-factor maps) and A-maps (local values of the logarithm of structure factor amplitudes at 15 Å) of EMD-8441 at different orientations.

### Immature prokaryote ribosomes

Next we processed immature ribosomal maps of the bacterial large subunit (Davis, Tan et al. 2016). These maps where obtained after depletion of bL17 ribosomal protein and are publicly available from the EMDB (EMD-8440, EMD-8441, EMD-8445, EMD-8450, EMD-8434) (Lawson, Baker et al. 2011). In this case, we focussed on showing the capacity of our proposed local occupancy maps to interpret and analyse reconstructions showing a high degree of compositional heterogeneity.

Figure 2 shows the obtained results. The first row shows the different maps to be processed as deposited in the EMDB. Next, we show the obtained occupancy maps, where the mature 50S ribosome (EMDB-8434) is coloured according to the corresponding occupancy maps. These figures clearly show regions that are lacking in the different immature maps with respect to the mature map. Thus, occupancy maps were used to create binary masks to segment the mature 50S ribosome map, extracting after the densities that are missing in the respective immature maps. These densities are shown in the third column of Figure 2 with different colours (yellow, red, indigo and green). The obtained occupancy maps also allow us to define a “maturity level” index. This index is calculated by comparing the number of voxels activated in the solvent mask of the mature 50S reconstruction with the ones in the occupancy masks (see methods section for a more detailed description). As can be seen from Figure 2, the larger the unfolded regions in the immature maps are, the smaller the maturity level is. This maturity level index allows us to quantitatively sort the different immature maps in a spectrum according to their maturity.

We further show the advantages of our LocSpiral and LocBFactor approaches in these highly heterogenous datasets. In the second row of Figure 2, we show maps with improved contrast at high-resolution obtained after processing EMD-8441 by the LocSpiral method and by Relion (Fernandez, Luque et al. 2008, Kimanius, Forsberg et al. 2016). The same soft mask was applied to both maps. In the figure, we show the maps at low and high threshold values. When a low threshold value is used, it is not possible to see details in the Relion map, while at high threshold values many regions of this map are not visible. Conversely, our LocSpiral approach shows high resolution features at both high and low thresholds without losing appreciable map densities.

Finally, we also show the obtained local B-factor map (B map) and the local values of the logarithm of structure factor’s amplitudes at 15 Å (A map in the figure). The average value of the local B-factor map to be used for sharpening is −299.11 Å^2^ (slope of the local Guinier plot multiplied by 4). We obtained the B-factor estimations within a resolution range between 15 Å to the FSC resolution given by 3.7 Å. Interestingly, the B-factor map shows lower B-factors in the outer part of the macromolecule, corresponding to regions that are folded partially and show compositional and conformational heterogeneity. As we will see in more detail in the next section, these modestly sloped Guinier plots come from low amplitudes (close or below to the noise amplitude level) at resolutions of 15 Å or higher. This result can be directly observed in the obtained A map in the figure that shows low values in potentially unfolded regions. Therefore, in order to accurately analyze local B-factor maps, it is necessary to interpret both B and A maps.

### Pre-catalytic spliceosome

Next we processed the Saccharomyces cerevisiae pre-catalytic B complex spliceosomal single particles deposited in EMPIAR (EMPIAR 10180) (Iudin, Korir et al. 2016, Plaschka, Lin et al. 2017). Because this dataset exhibits a high degree of conformational heterogeneity, we concentrate here on showing the capacity of our LocBFactor method. We used the approach described in (Gomez-Blanco, Kaur et al. 2019) to obtain a reconstruction at 4.28 Å resolution after Relion postprocessing (Fernandez, Luque et al. 2008, Kimanius, Forsberg et al. 2016). Then, the unfiltered map provided by Relion autorefine was used to test our proposed methods.

In Figure 3A, we show a central slice along the Z axis of this map, and several points are marked with coloured squares. These points show parts of the map that correspond to clear spliceosome densities (green and red), flexible and low resolution spliceosomal regions (yellow and blue) and background (magenta). Figure 3B shows the corresponding Guinier plots of these points. Solid lines represent measured values of the logarithm of SNR-weighted structure factor amplitudes, while dashed lines show fitted curves. This figure also provides the obtained B-factors for the different curves. The Guinier plots and B-factors are determined within a resolution range of 15 Å to the FSC resolution, given by 4.28 Å. As can be seen from Figure 3B, the red and green curves, which correspond to clear spliceosomal densities, present high amplitude values at 15 Å, while yellow, blue and magenta curves show low amplitudes at 15 Å and a flat profile within the resolution range. In Figure 3B we also show in the black curve, the Guinier plot of the noise/background amplitudes obtained from the 95% quantile of the empirical noise/background distribution for reference. The discontinuous black line indicates the linear fit of this noise Guinier plot. Comparing the yellow, blue and magenta curves, it is clear that these plots are below our noise level and that the shape of these curves is similar to that of the noise curve. Figure 3D shows the spliceosome map coloured with the obtained B-factor map to be used for sharpening (slope of the local Guinier plot multiplied by 4). The average of these B-factors for sharpening is −153.68 Å^2^, while the value reported by Relion postprocess is −158.08 Å^2^. Figure 3E shows a central slice of the obtained B-factor map along Z axis. As can be seen from Figure 3E, the B-factor values in the background are low since the corresponding Guinier plots show a flat spectrum within the resolution range. In Figure 3F, we show the local values of the logarithm of structure factor’s amplitudes at 15 Å (A map). As expected, this map shows low amplitudes at highly flexible and moving regions. These results show that B-factors calculated from the EM map present modestly sloped Guinier plots in very flexible and blurred regions, which is not in agreement with the concept of B-factor as a measure of position uncertainty or disorder. The explanation for this discrepancy lays in the resolution range used to calculate the local B-factors from the electron microscopy map. Within the resolution range of [15, 4.28] Å, the amplitudes of the highly flexible parts (helicase and SF3b domains) are very low and below the noise level, showing a flat spectrum that can be seen in Figure 3E. Thus, these regions show very low signal-to-noise rations and are just noise within this resolution range. Therefore, we have recalculated our B-factors using a new resolution range of [20, 10] Å. The results are shown in Figure S2. As can be seen from Figure S2, now the flexible parts show low B-factor and amplitudes at 20 Å^2^. In this case, the average value of the local B-factor map to be used for sharpening is of −639.15 Å^2^ (slope of the local Guinier plot multiplied by 4).

**Figure 3.**
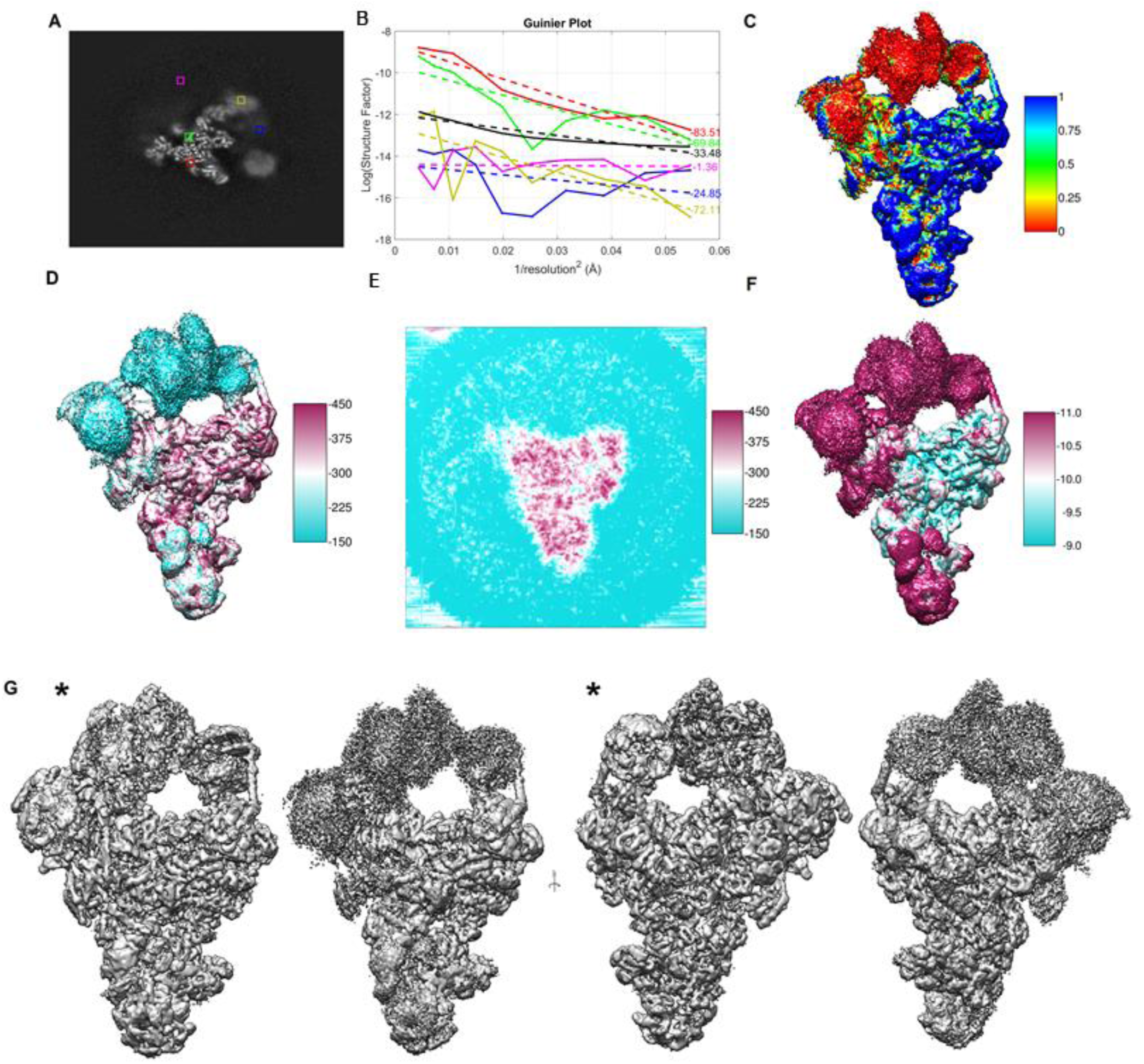
A) Central slice along Z axis of obtained Saccharomyces cerevisiae pre-catalytic B complex spliceosome map using EMPIAR 10180 single particles. Coloured squares mark parts of the map corresponding to clear spliceosome densities (green and red), flexible and low resolution spliceosomal regions (yellow and blue) and background (magenta); B) Guinier plots at map points indicated in the coloured squares. Solid lines represent SNR-weighted values of the logarithm of structure factor amplitudes, while discontinued lines show the fitted lines; C) Spliceosome map coloured with the obtained occupancy map; D) and E) Spliceosome map and spliceosome central slice coloured with the obtained B-factor map; F) Spliceosome map coloured with the obtained A map; G) Maps at different orientations and similar threshold values obtained by our local enhancement method and by the postprocessing method of Relion. The map obtained by our proposed method is marked with an asterisk.

We also show results obtained by the LocOccupancy and LocSpiral methods for this highly heterogenous case. Figure 3C show the spliceosome map coloured according to the occupancy map. From Figure 3C, we see that the flexible and moving parts of the spliceosome show low occupancies. Finally, in Figure 3G, we show maps at different orientations and similar threshold values obtained by our LocSpiral and by the postprocessing method of Relion 3 (Fernandez, Luque et al. 2008, Kimanius, Forsberg et al. 2016). The map obtained by our proposed method is marked with an asterisk. As before, the map obtained by our proposed approach shows fewer fragmented and broken densities, especially in the flexible part of the spliceosome reconstruction, and enhanced details in the central core portion.

### SARS-CoV-2

We have processed recent cryo-EM maps of the CoV spike (S) glycoprotein (Walls, Park et al. 2020, Wrapp, Wang et al. 2020). These maps include cryo-EM reconstructions of the SARS-CoV-2 spike in the prefusion conformation with a single receptor-binding domain (RBD) up (EMD-21375) and after imposing C3 symmetry in the refinement to improve visualization of the symmetric S2 subunit (EMD-21374). We also processed additional cryo-EM reconstructions from the Veesler lab of the SARS-CoV-2 spike glycoprotein with three RBDs down (EMD-21452) and the SARS-CoV-2 spike ectodomain structure (EMD-21457) with a single RBD up. The reported global resolution of these maps is 3.46 Å, 3.17 Å, 2.8 Å and 3.2 Å, respectively. Interesting, deposited atomic models (PDBs PDB 6vsb, PDB 6vxx and PDB 6vyb) incompletely cover the reconstructed cryo-EM maps, showing the existence of disordered or over sharpened regions after B-factor correction that could not be modelled. Figure S3 displays corresponding maps and fitted atomic models showing the large amount of protein that is not currently modelled.

In Figure 4A, we show deposited EMD and obtained maps by our proposed LocSpiral approach. In this figure, we use a relatively low threshold to visualize the outer parts of the protein. This figure shows that our obtained reconstructions present less fragmented and broken densities and better map connectivity than the deposited EMD maps, suggesting that our approach improves the analysis and visualization of the outer regions and potentially aides in the modelling of additional map motifs. Interesting, the obtained EMD-21374 map shows some fragmented density at the top of the spike, however, we believe that this additional density is an artifact that comes as result of artificially imposing C3 symmetry on particles that are asymmetric. Then, we used the reconstruction obtained from LocSpiral of EMD-21375 and to improve the deposited atomic model (PDB 6vsb). As result, we could model additional loops and motifs: K444.C-F490.C; E96.C-S98.C; NAG1322.C; P812.C-K814.C, and some additional amino acids, which are now visible in the improved map: P621.C-G639.C; S673.C-V687.C; A829.B-A825.B. We were also able to visualize density corresponding to numerous additional *N*-linked glycans that could not be resolved in the original reconstruction. Examples of some regions that could be further modelled are shown in Figures 4C and 4D. In Figure 4C, we show at the left the obtained LocSpiral map with the improved atomic model in green, and at the right the deposited EMD map with the PDB 6vsb in magenta. Figure 4D shows in white the PDB 6vsb with the traced parts of the glycan proteins marked with purple spheres and in red the additional traced parts using our improved LocSpiral map. In addition, in this figure we provide zoomed views of two glycan proteins that could be further modelled with our improved map also shown in the image.

**Figure 4.**
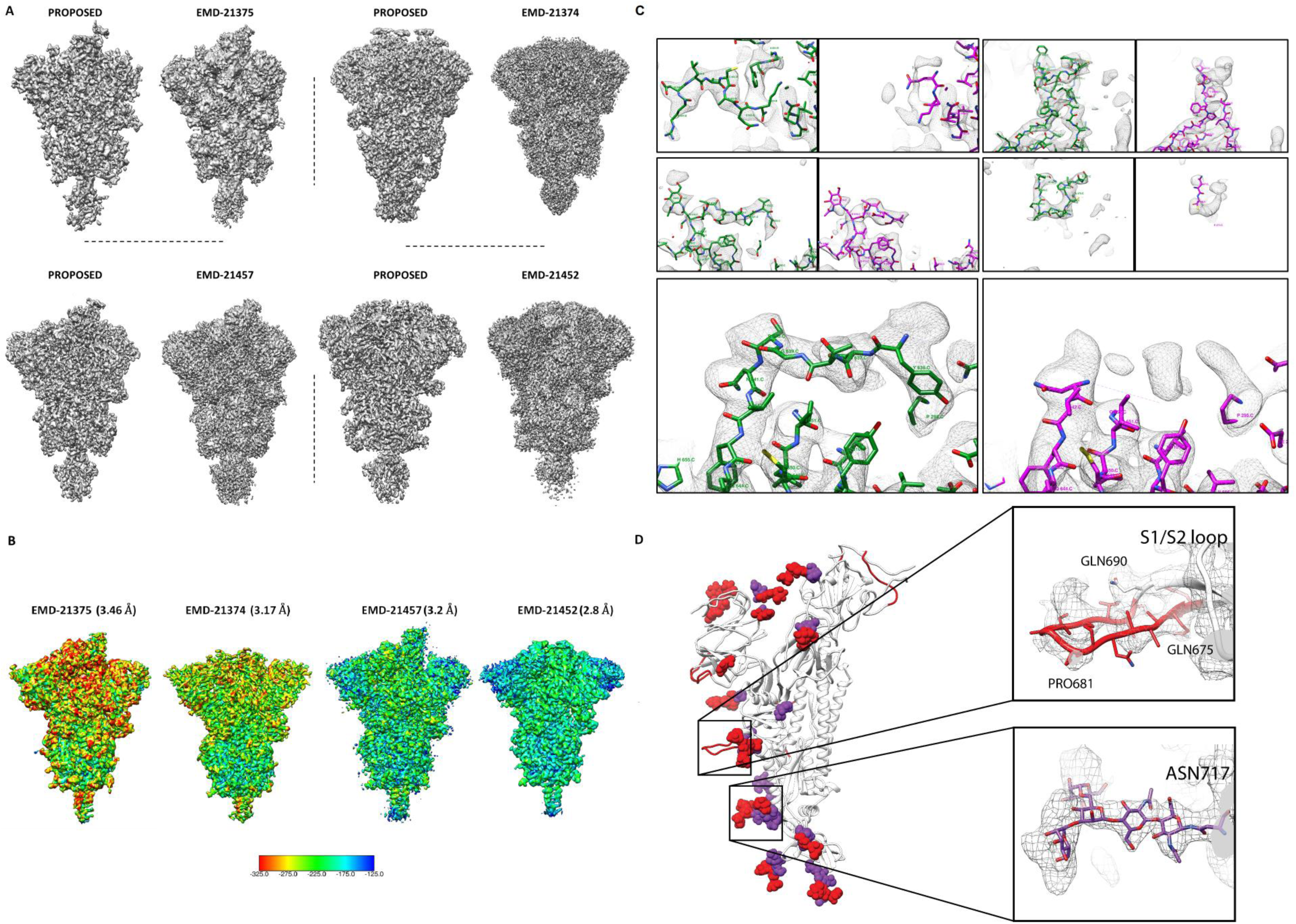
A) Maps obtained by our proposed LocScale approach (left) and deposited maps in EMDB with accessing codes (EMD-21375, EMD-21374, EMD-21457, EMD-21452); B) Obtained B-factor maps for EMD-21375, EMD-21374, EMD-21457, EMD-21452; C) Visual examples of some regions that could be further modelled after processing EMD-21375 with LocSpiral, where at the left are shown the LocSpiral maps with the improved atomic models in green, and at the right the deposited EMD-21375 map with the PDB 6vsb in magenta; D) In white, PDB 6vsb with traced parts of the glycan proteins marked with purple spheres. In red, additional parts that could be traced using our improved LocSpiral map. Inside the black squares, zoomed views of two glycan proteins that could be further modelled.

Finally, in Figure 4B, we show obtained local B-factor maps using a similar colormap and estimated global resolution estimations. These figures show that EMD-21452 and EMD-21454 show lower B-factors than EMD-21375 and EMD-21374, and then a better localizability of secondary structure and residues.

### Apoferritin

We have also applied these techniques to recently reported high-resolution cryo-EM reconstructions of mouse apoferritin (EMD-9865) and (EMD-21024). The reported global resolution of these reconstructions are 1.54 and 1.75 Å for EMD-9865 and EMD-21024, respectively.

In Figure 5, we show the results of our obtained B-factor map to be used for sharpening (slope of the local Guinier plot multiplied by 4), local amplitudes at 15 Å and local occupancy maps. The resolution range used to estimate B and A maps was between 15 Å to the reported global resolution for both cases. The occupancy maps were calculated for these high-resolution maps between 5 to the global resolution. As can be seen from Figure 5, EMD-9865 shows lower B-factors and higher local amplitudes, indicating a better-quality reconstruction. In both cases, the highest B-factors are in the outer regions of the protein. Additionally, local occupancies show similar maps for both cases, showing occupancies as low as approximately 0.5 at the outer part and indicating the presence of flexibility in these outer residues. Note that the obtained average and standard deviation of B-factors inside a solvent mask: −56 Å^2^ and 7.20 Å^2^ (EMD-9865) and - 78 Å^2^ and 8.93 Å^2^ (EMD-21024) respectively, which reflects the high quality of these reconstructions.

**Figure 5.**
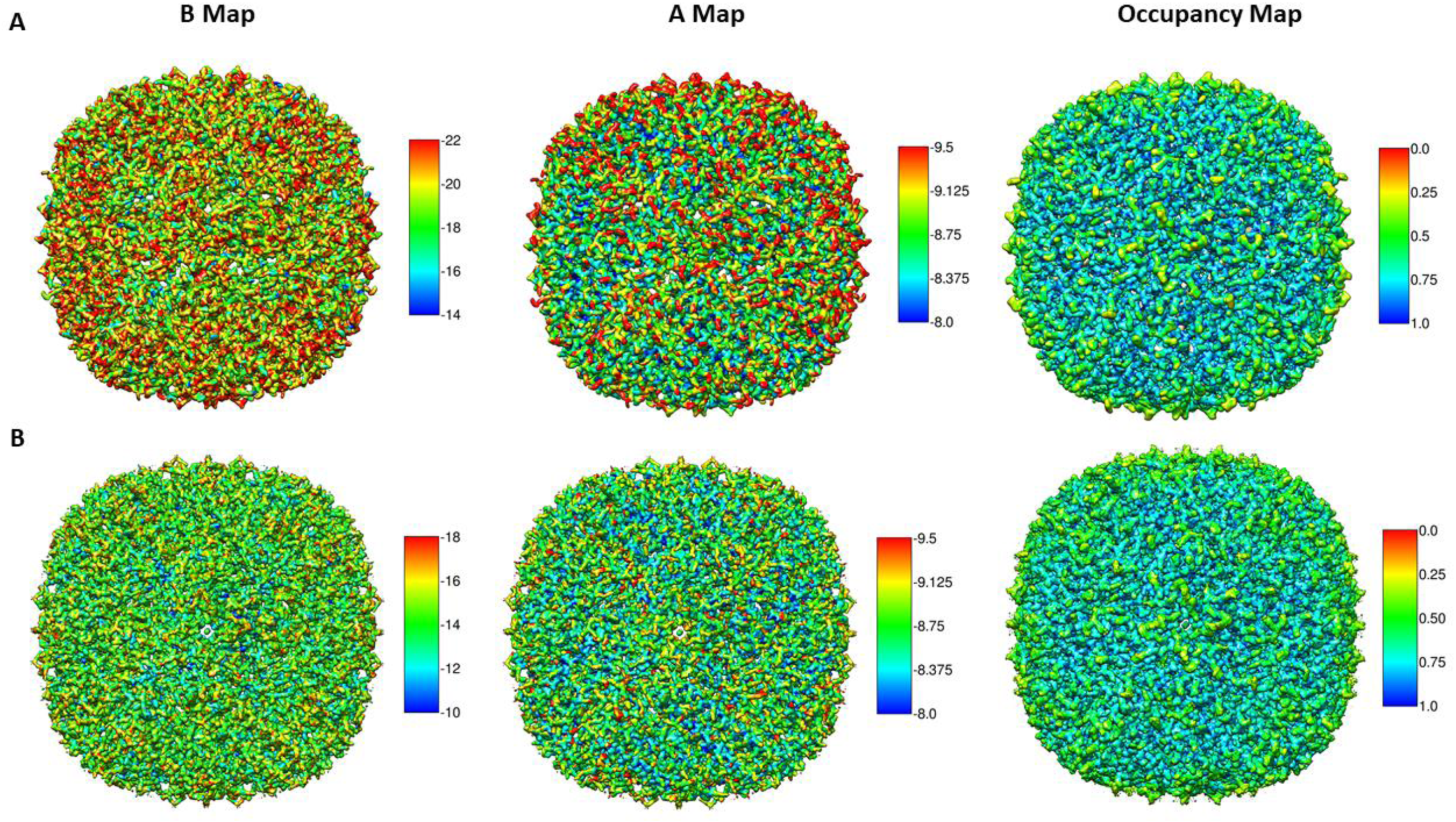
Obtained B-factor, A and occupancy maps for EMD-21024 (A) and EMD-9865 (B)

## Discussion

In this paper, we have introduced methods to improve the analysis and interpretability of cryo-EM maps. These methods include map enhancement approaches (LocSpiral and local B-factor sharpening), and approaches to calculate local B-factors and density occupancy maps. We have shown in our experiments that our LocSpiral approach improves map connectivity showing fewer fragmented and broken densities and better coverage of the atomic model. In fact, our LocSpiral approach has been applied on several published publications (Ichikawa, Khalifa et al. 2019, Yang, Chen et al. 2019, Gutmann, Schafer et al. 2020, Jahagirdar, Jha et al. 2020, Khalifa, Ichikawa et al. 2020), enabling molecular modelling on maps with flexibility and light anisotropic resolution. We envision that our proposed methods to estimate local B-factors and occupancy maps could be used to improve *de novo* model building. First, these maps can be employed to guide the manual tracing. These maps can be informative to estimate the range of structures that could be compatible to the given electron microscopy density. Second, for very high resolution cryo-EM maps, these values can be used as an approximation of the atomic B-factors and occupancies to be further refined as part of the automatic model refinement process by automatic model building packages as Phenix (Liebschner, Afonine et al. 2019) or Refmac (Winn, Murshudov et al. 2003). B-factor maps provide complementary information to local resolution maps, though usually these results are correlated. The latter usually determines the resolution at a given point by comparing the map to noise or background amplitudes (Vilas, Gomez-Blanco et al. 2018), while the former determines the rate of signal amplitude fall off within a resolution range. We can find map regions with similar local resolution (map amplitude similar to noise/background amplitude at this resolution and coordinates), while different B-factor as the signal damping could be different within the used resolution range (highly or slowly sloped).

We have seen that we have to be careful when processing maps affected by high flexibility and heterogeneity as the obtained B-factors could be underestimated if the selected resolution range is above the local resolution at these regions. However, these problematic cases can be easily detected as the A map values in these locations are close to or below the noise amplitude. Thus, these regions can be automatically disable and not taken into consideration.

The methods proposed here are semi-automated and essentially only require the map to enhance or analyse, a binary solvent mask and a resolution range as inputs. They do not require additional information as atomic models or local resolution maps. The common link between all these approaches is the use of the spiral phase transform, which is used to factorize cryo-EM maps into amplitude and phase terms in real space for different resolutions. The spiral phase transform has been extensively used in optics for phase extraction in interferometry (Larkin, Bone et al. 2001, Antonio Quiroga and Servin 2003, Vargas, Quiroga et al. 2011, Vargas, Restrepo et al. 2011, Vargas, Quiroga et al. 2012) or by Shack-Hartmann sensors (Vargas, González-Fernandez et al. 2010, Vargas, Restrepo et al. 2012). This transformation is not new in cryo-EM as it has been proposed previously to facilitate particle screening (Vargas, Abrishami et al. 2013), CTF estimation (Vargas, Oton et al. 2013) and local and directional resolution determination (Vilas, Gomez-Blanco et al. 2018, Vilas, Tagare et al. 2020). In (Vilas, Gomez-Blanco et al. 2018, Vilas, Tagare et al. 2020) the authors used the Riesz transform to obtain amplitude maps, which is similar to the spiral phase transform.

Cryo-EM reconstructions of different types of macromolecules have been used to test the performance of these algorithms. Specifically, we have used a membrane protein (TRP channel), immature ribosomes affected by high compositional heterogeneity, the spliceosome that shows high conformational heterogeneity, recent SARS-CoV-2 reconstructions exhibiting dynamic regions and high resolution apoferritin reconstructions. In all cases, our proposed approaches show excellent results, improving the analysis and the interpretability of the processed maps. The proposed methods are also highly efficient. For example, the processing of EMD-21457 (map size 400 px^3^) using our local enhancement approach took only 12 min on a standard laptop using 4 cores.

## Acknowledgments

This work was supported by grants from NSERC Discovery Grant (RGPIN-2018-04813), J.V. acknowledges economical support from the Ramón y Cajal 2018 program (RYC2018-024087-I). We want to thank helpful discussions with Jose Jesus Fernandez.

## Methods

The proposed methods are based on a 3D generalization of the 2D spiral phase transform. In the following, we present the 3D spiral phase transform and its application to map enhancement, local B-factor determination, and estimation of local map occupancies.

### 3D spiral phase transform

The spiral phase transform is a Fourier operator that can factorize a 3D map into its amplitude and phase terms in real space at different resolutions. We assume without loss of generality that a given 3D map can be modelled as a 3D phase modulated signal given by

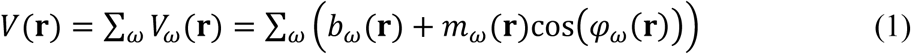

where *V* is the <u>cryo</u>-EM map, *V*_*ω*_ is a band-passed map filtered at frequency *ω, b*_*ω*_ the 3D background or DC term, *m*_*ω*_ the 3D amplitude map, φ_*ω*_ the 3D modulating phase and r = (*x, y, z*). Assuming that we are interested in spatial frequencies higher than 1/50-1/30 1/Å and that the background is usually a low frequency signal, we can approximate a high-passed filtered map *V*_*HP*_ for resolutions higher than 50-30 Å as

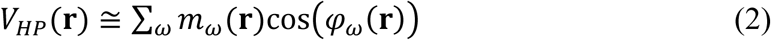

The quadrature transformation of Eq. (2) is given by

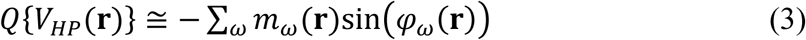

Assuming that *m*_*ω*_ is a low varying map compared to *φ*_*ω*_, the gradient of *V*_*HP*_ is approximated by

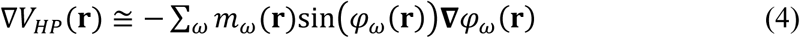

Rearranging terms, we obtain

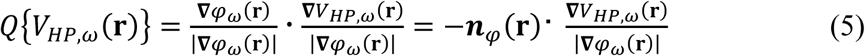

Eq. (5) shows that the quadrature term is composed by two terms. The first is an orientation map *n*_*φ*_ and the second corresponds to a non-linear operator that can be interpreted as a 3D generalization of the 1D Hilbert transform, which can be efficiently calculated using the Fourier transform. As shown in (Servin, Quiroga et al. 2003), the operator **∇***V*_*HP,ω*_(**r**)/|**∇***φ*_*ω*_(**r**)| corresponds to the 3D Hilbert transform for our band-passed maps *V*_*HP,ω*_(**r**) then

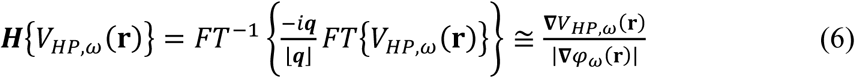

Thus, Eq. (5) can be rewritten as

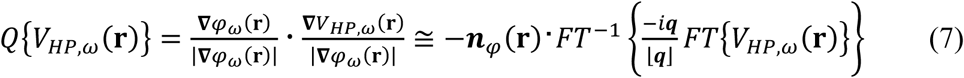

Note that *n*_*φ*_ is an unit vector pointing in the same direction that **∇***V*_*HP,ω*_(**r**) (remember that *m*_*ω*_ is a low varying map compared to *φ*_*ω*_), but maybe with different orientation because a possible change of sign introduced by the cosine term in Eq. (2). We can rewrite Eq. (7) as

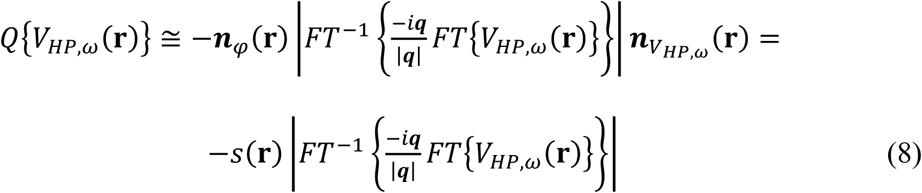

where *s*(**r**) is a function with range +1 or −1 considering that *n*_*φ*_(**r**) and 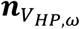 can be parallel or antiparallel. From Eq. (8), we can obtain an estimation of *φ*_*ω*_(**r**) affected by an indetermination in its sign

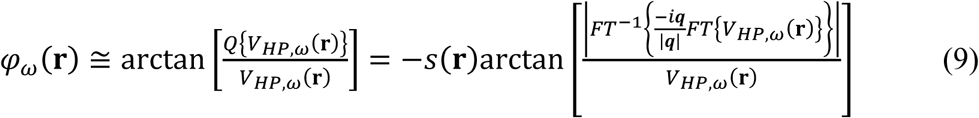

However, we can use Eq. (9) to obtain the modulation and cosine terms in Eq. (2) separately without sign ambiguity as

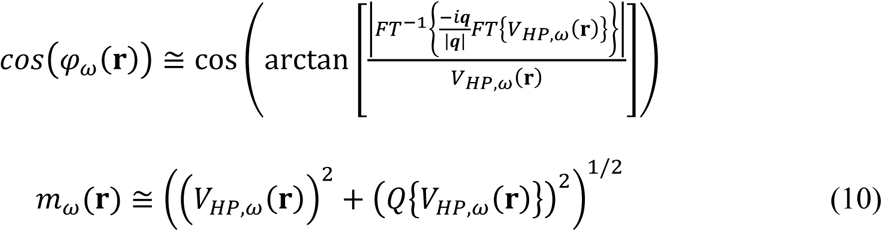

Using these expressions, we can obtain for each frequency *ω* the terms cos(*φ*_*ω*_(**r**)) and *m*_*ω*_(**r**).

### Local enhanced map (LocSpiral)

We are proposing here a robust local map enhancement method that only requires as input a binary mask of the macromolecule. The approach works for both high and moderate resolution maps. In the following, we provide details of the proposed method.

As explained before, each band-pass filtered map can be factorized into an amplitude and phase term by the spiral phase transform. Then, given a user defined solvent mask, the method obtains the empirical noise amplitude probability distribution 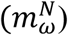 at frequency *ω*, selecting the voxels not included in the solvent mask. From this distribution, the approach determines the noise amplitude value corresponding to the 95% quantile, given by 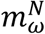(*q*=95%). This value is used to locally normalize map amplitudes in real space along different frequencies and remove local signals that are below this amplitude threshold as they are likely noise at this given frequency and position. After this nonlinear amplitude transformation, the map is given by

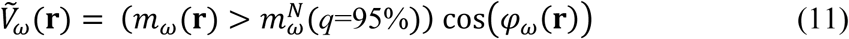

Then, Eq. (2) is rewritten as

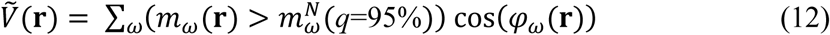

The method allows as option the use of a SNR weighting parameter to weight the contribution of the different amplitudes in the final map. In this case, Eq. (12) is rewritten as

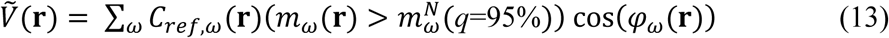

with *C*_*ref,ω*_(**r**) the SNR weighting parameter given by

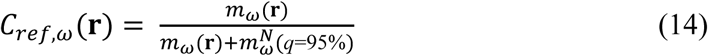

### Local B-factor determination (LocBFactor)

The factorization of a 3D map into its amplitude and phase terms in real space for different resolutions allows the efficient determination of local B-factor maps. For resolutions between 15-10 Å to the estimated global map resolution, the method first obtains the local amplitude maps *m*_*ω*_(**r**). These amplitude maps are then used to obtain SNR-weighted log-amplitudes of the structure factors locally as

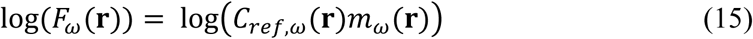

with *C*_*ref,ω*_(**r**) a SNR weighting parameter defined in (14). These expressions can be used to fit log(*F*_*ω*_(**r**)) versus *ω*^2^ within a resolution rage between 15-10 Å to the estimated global map resolution. Thus, finally we have

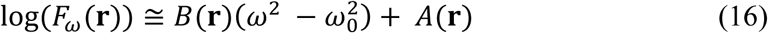

with *B*(**r**) the local B-factor map and *A*(**r**) the log-amplitude map intensities at *ω*_0_, which corresponds to the lowest frequency within the used frequency range.

### Local B-factor sharpened map (LocBSharpen)

The spiral phase transform can be used to obtain local B-factor sharpened maps. Note that Expression (2) can be modified for frequencies higher than *ω*_0_ as

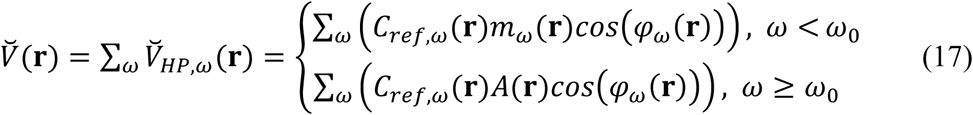

### Local occupancy map (LocOccupancy)

Low occupancy map regions correspond to parts of the macromolecule where map amplitudes of the reconstruction are significantly smaller when compared to other regions of the macromolecule. Keeping this in mind, we define the occupancy map as

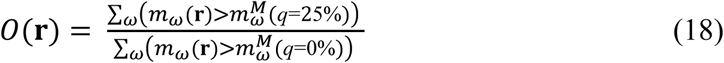

where 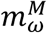(*q*=25%) is obtained from the empirical macromolecule amplitude probability distribution 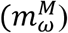 at frequency *ω*. This amplitude probability distribution is calculated from voxels that are included in the solvent mask. From this distribution, the approach determines the macromolecule amplitude values corresponding to the 25% and 0% quantiles, given by 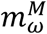(*q*=25%) and 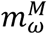(*q*=0%) that are used as threshold. To calculate local occupancy maps, a typical resolution range between 50 and 10-8 Å is used to obtain density occupancies of complete secondary structure motifs, while ranges between 5 to 3-1.5 Å are used for high resolution cryo-EM maps to obtain occupancies of residues.

### Maturity level index

In the analysis of the immature 50S ribosomes, we have proposed a maturity level index. This index can be extended to the analysis of any maturing macromolecule and is useful to place immature macromolecules into a maturing timeline. The calculation of this index requires reconstructions of immature and mature macromolecules. The mature reconstruction is used to obtain a binary solvent mask, while the immature reconstructions are used to calculate occupancy maps. These occupancy maps allow us to determine highly occupied regions (occupancy >0.75) and calculate occupancy masks. Then, the index is obtained comparing the number of voxels activated in the solvent mask of the mature reconstruction with the ones in the occupancy masks. As can be seen from Figure 2, the larger are the regions that are not folded in the immature maps, the smaller is the maturity level.

## Software and data availability

The source code for the presented methods are freely available under the terms of an open source software license and can be downloaded from https://github.com/1aviervargas/LocSpiral-LocBSharpen-LocBFactor-LocOccupancy. Previously published datasets used for testing are available from the Electron Microscopy Data Bank (https://www.ebi.ac.uk/pdbe/emdb/) under accession codes EMD-10418, EMD-8440, EMD-8441, EMD-8445, EMD-8450, EMD-8434, EMD-21375, EMD-21374, EMD-21452 and EMD-21457. Resultant maps by our proposed approaches can be downloaded from https://github.com/1aviervargas/LocSpiral-LocBSharpen-LocBFactor-LocOccupancy.

## Author contributions

J.V had the idea, J.V, S.K and J.G and devised the theory, developed and implemented the algorithm, performed the experiments and wrote the manuscript. R.S.G helped analysing and interpreting data. A.K, D. W., J.S.M and H.B analysed data, wrote part of the manuscript and provided comments and feedback. All authors reviewed the manuscript, supervised the experiments and discussed the results.

## Supplementary Material

**Figure S1.**
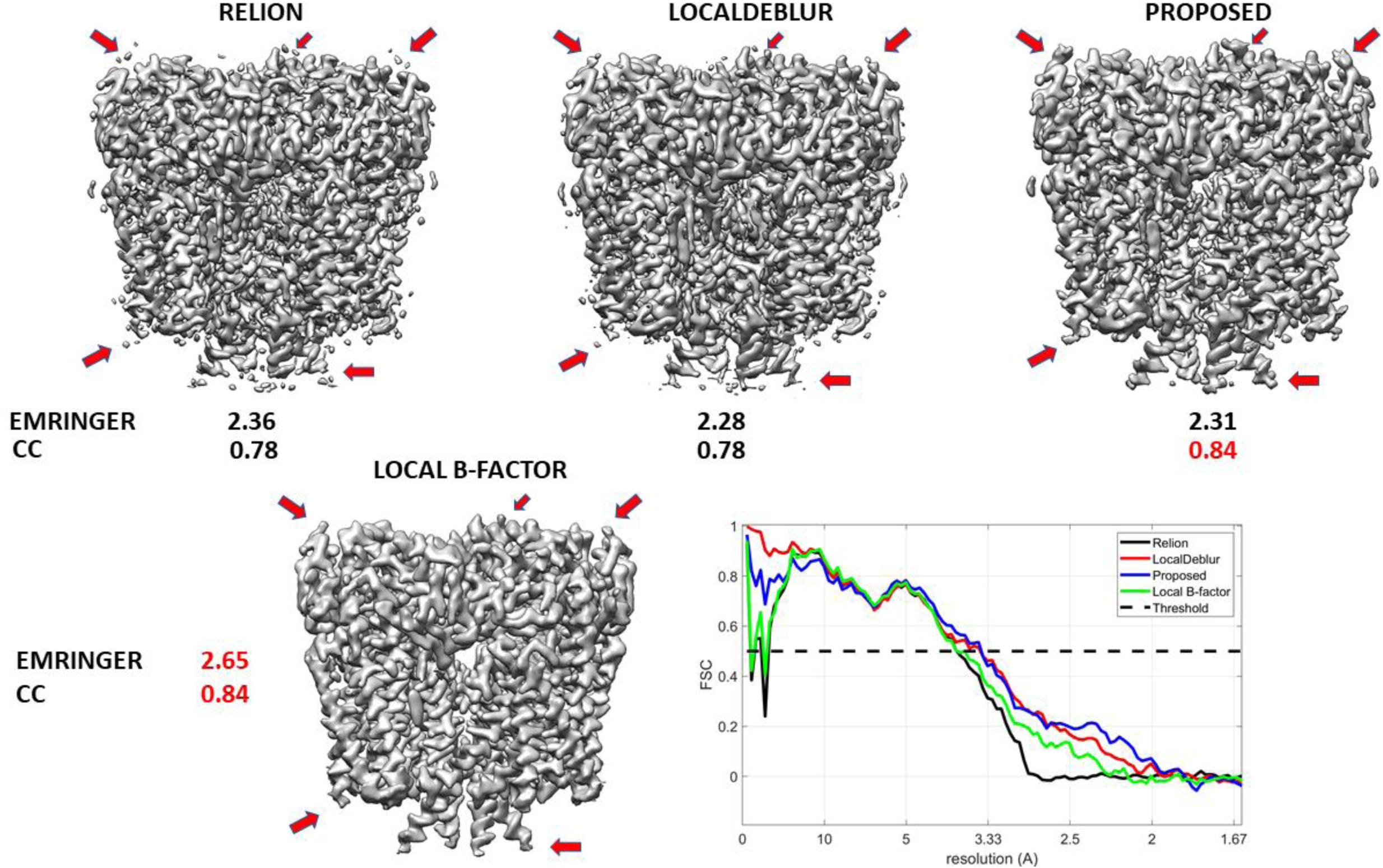
Comparison between maps obtained with the global B-factor correction approach as implemented in Relion, LocalDeblur, our proposed map enhancement approach and our proposed local B-factor correction method. Red arrows show broken or missed densities that are shown in our obtained maps. Below each map EMRINGER and cross-correlation (CC) scores calculated between obtained maps and the atomic model (PDB 6t9n) are provided. We also show FSC curves comparing the different maps with the reference atomic model (PDB 6t9n).

**Figure S2.**
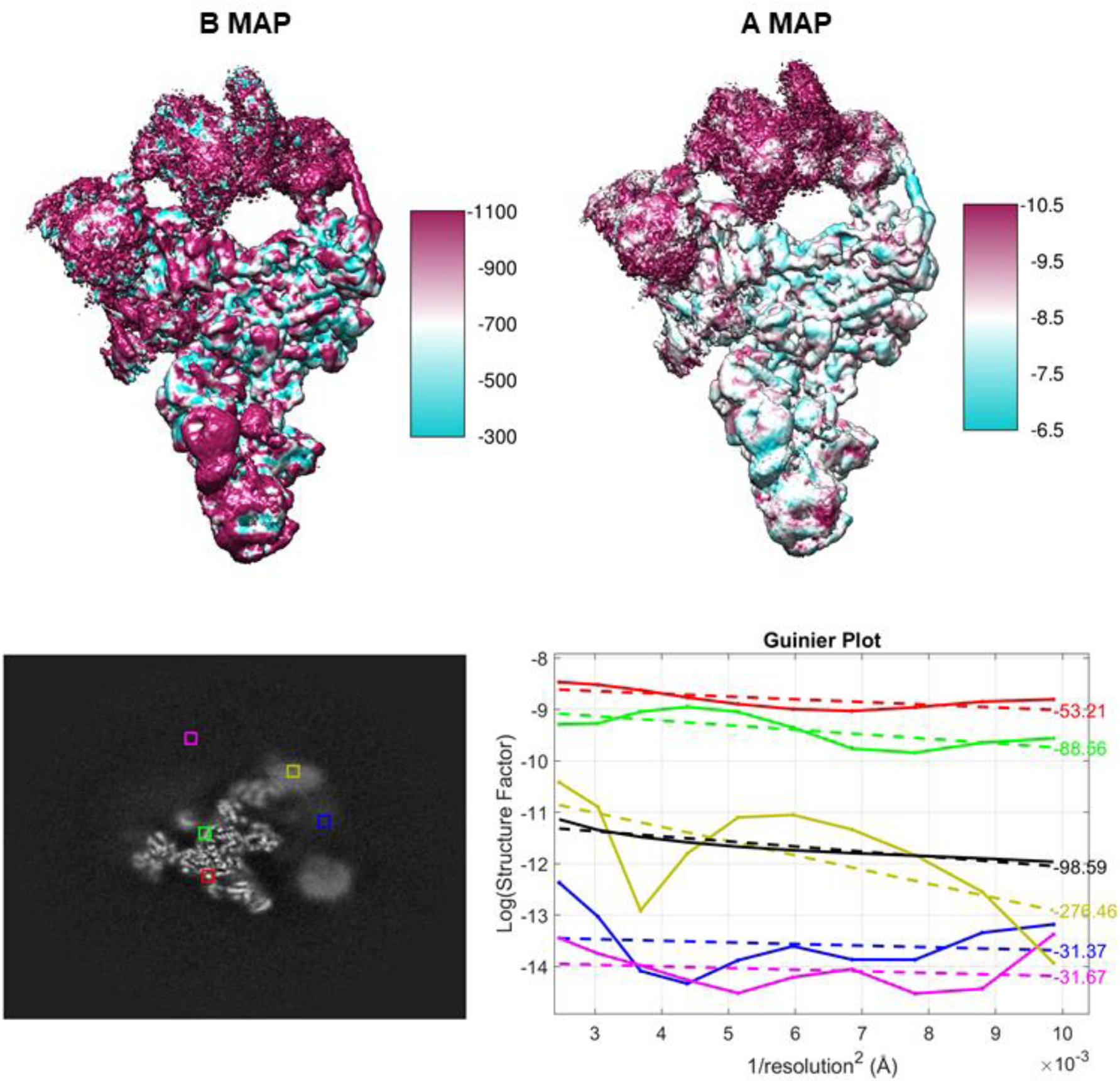
Improved maps by our proposed enhancement approach obtained from EMD-21375, EMD-21457, EMD-21452 and corresponding fitted atomic models.

**Figure S3.**
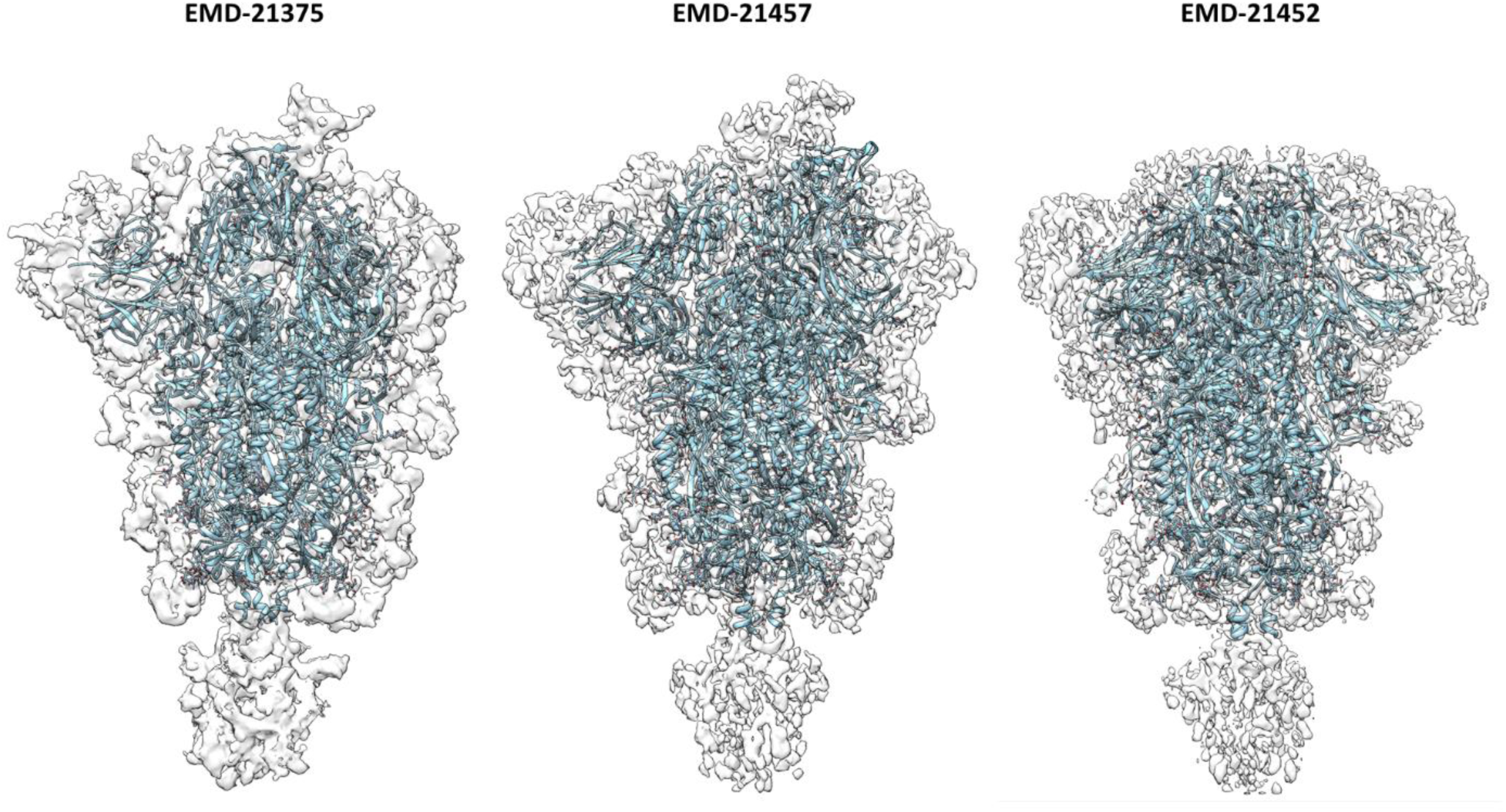

## References

Afonine, P. V., B. P. Klaholz, N. W. Moriarty, B. K. Poon, O. V. Sobolev, T. C. Terwilliger, P. D. Adams and A. Urzhumtsev (2018). “New tools for the analysis and validation of cryo-EM maps and atomic models.” Acta Crystallogr D Struct Biol 74(Pt 9): 814–840.

Antonio Quiroga, J. and M. Servin (2003). “Isotropic n-dimensional fringe pattern normalization.” Optics Communications 224(4): 221–227.

Barad, B. A., N. Echols, R. Y. Wang, Y. Cheng, F. DiMaio, P. D. Adams and J. S. Fraser (2015). “EMRinger: side chain-directed model and map validation for 3D cryo-electron microscopy.” Nat Methods 12(10): 943–946.

Davis, J. H., Y. Z. Tan, B. Carragher, C. S. Potter, D. Lyumkis and J. R. Williamson (2016). “Modular Assembly of the Bacterial Large Ribosomal Subunit.” Cell 167(6): 1610–1622 e1615.

Erney Ramirez-Aportela, J. L. V., Roberto Melero, Pablo Conesa, Marta Martinez, David Maluenda, Javier Mota, Amaya Jimenez, Javier Vargas, Roberto Marabini, Jose Maria Carazo, Carlos Oscar Sanchez Sorzano (2017). “Automatic local resolution-based sharpening of cryo-EM maps.” biorxiv.

Fernandez, J. J., D. Luque, J. R. Caston and J. L. Carrascosa (2008). “Sharpening high resolution information in single particle electron cryomicroscopy.” J Struct Biol 164(1): 170–175.

Ge, P., D. Scholl, N. S. Prokhorov, J. Avaylon, M. M. Shneider, C. Browning, S. A. Buth, M. Plattner, U. Chakraborty, K. Ding, P. G. Leiman, J. F. Miller and Z. H. Zhou (2020). “Action of a minimal contractile bactericidal nanomachine.” Nature.

Gomez-Blanco, J., S. Kaur, J. Ortega and J. Vargas (2019). “A robust approach to ab initio cryoelectron microscopy initial volume determination.” J Struct Biol 208(3): 107397.

Gutmann, T., I. B. Schafer, C. Poojari, B. Brankatschk, I. Vattulainen, M. Strauss and U. Coskun (2020). “Cryo-EM structure of the complete and ligand-saturated insulin receptor ectodomain.” J Cell Biol 219(1).

Ichikawa, M., A. A. Z. Khalifa, S. Kubo, D. Dai, K. Basu, M. A. F. Maghrebi, J. Vargas and K. H. Bui (2019). “Tubulin lattice in cilia is in a stressed form regulated by microtubule inner proteins.” 201911119.

Iudin, A., P. K. Korir, J. Salavert-Torres, G. J. Kleywegt and A. Patwardhan (2016). “EMPIAR: a public archive for raw electron microscopy image data.” Nat Methods 13(5): 387–388.

Jahagirdar, D., V. Jha, K. Basu, J. Gomez-Blanco, J. Vargas and J. Ortega (2020). “Alternative Conformations and Motions Adopted by 30S Ribosomal Subunits Visualized by Cryo-Electron Microscopy.” bioRxiv: 2020.2003.2021.001677.

Jakobi, A. J., M. Wilmanns and C. Sachse (2017). “Model-based local density sharpening of cryo-EM maps.” Elife 6.

Khalifa, A. A. Z., M. Ichikawa, D. Dai, S. Kubo, C. S. Black, K. Peri, T. S. McAlear, S. Veyron, S. K. Yang, J. Vargas, S. Bechstedt, J. F. Trempe and K. H. Bui (2020). “The inner junction complex of the cilia is an interaction hub that involves tubulin post-translational modifications.” Elife 9.

Kimanius, D., B. O. Forsberg, S. H. Scheres and E. Lindahl (2016). “Accelerated cryo-EM structure determination with parallelisation using GPUs in RELION-2.” Elife 5.

Larkin, K. G., D. J. Bone and M. A. Oldfield (2001). “Natural demodulation of two-dimensional fringe patterns. I. General background of the spiral phase quadrature transform.” Journal of the Optical Society of America A 18(8): 1862–1870.

Lawson, C. L., M. L. Baker, C. Best, C. Bi, M. Dougherty, P. Feng, G. van Ginkel, B. Devkota, I. Lagerstedt, S. J. Ludtke, R. H. Newman, T. J. Oldfield, I. Rees, G. Sahni, R. Sala, S. Velankar, J. Warren, J. D. Westbrook, K. Henrick, G. J. Kleywegt, H. M. Berman and W. Chiu (2011). “EMDataBank.org: unified data resource for CryoEM.” Nucleic Acids Res 39(Database issue): D456–464.

Liao, H. Y. and J. Frank (2010). “Definition and estimation of resolution in single-particle reconstructions.” Structure 18(7): 768–775.

Liebschner, D., P. V. Afonine, M. L. Baker, G. Bunkoczi, V. B. Chen, T. I. Croll, B. Hintze, L. W. Hung, S. Jain, A. J. McCoy, N. W. Moriarty, R. D. Oeffner, B. K. Poon, M. G. Prisant, R. J. Read, J. S. Richardson, D. C. Richardson, M. D. Sammito, O. V. Sobolev, D. H. Stockwell, T. C. Terwilliger, A. G. Urzhumtsev, L. L. Videau, C. J. Williams and P. D. Adams (2019). “Macromolecular structure determination using X-rays, neutrons and electrons: recent developments in Phenix.” Acta Crystallogr D Struct Biol 75(Pt 10): 861–877.

Murshudov, G. N. (2016). “Refinement of Atomic Structures Against cryo-EM Maps.” Methods Enzymol 579: 277–305.

Penczek, P. A. (2010). “Image restoration in cryo-electron microscopy.” Methods Enzymol 482: 35–72.

Plaschka, C., P. C. Lin and K. Nagai (2017). “Structure of a pre-catalytic spliceosome.” Nature 546(7660): 617–621.

Razi, A., J. H. Davis, Y. Hao, D. Jahagirdar, B. Thurlow, K. Basu, N. Jain, J. Gomez-Blanco, R. A. Britton, J. Vargas, A. Guarne, S. A. Woodson, J. R. Williamson and J. Ortega (2019). “Role of Era in assembly and homeostasis of the ribosomal small subunit.” Nucleic Acids Res.

Rosenthal, P. B. and R. Henderson (2003). “Optimal determination of particle orientation, absolute hand, and contrast loss in single-particle electron cryomicroscopy.” J Mol Biol 333(4): 721–745.

Scheres, S. H. (2015). “Semi-automated selection of cryo-EM particles in RELION-1.3.” J Struct Biol 189(2): 114–122.

Servin, M., J. A. Quiroga and J. L. Marroquin (2003). “General n-dimensional quadrature transform and its application to interferogram demodulation.” Journal of the Optical Society of America A 20(5): 925–934.

Sherwood, D., J. Cooper and D. Sherwood (2011). Crystals, x-rays, and proteins : comprehensive protein crystallography. New York, Oxford University Press.

Terwilliger, T. C., O. V. Sobolev, P. V. Afonine and P. D. Adams (2018). “Automated map sharpening by maximization of detail and connectivity.” Acta Crystallogr D Struct Biol 74(Pt 6): 545–559.

Vargas, J., V. Abrishami, R. Marabini, J. M. de la Rosa-Trevin, A. Zaldivar, J. M. Carazo and C. O. S. Sorzano (2013). “Particle quality assessment and sorting for automatic and semiautomatic particle-picking techniques.” J Struct Biol 183(3): 342–353.

Vargas, J., L. González-Fernandez, Juan A. Quiroga and T. Belenguer (2010). “Shack–Hartmann centroid detection method based on high dynamic range imaging and normalization techniques.” Applied Optics 49(13): 2409–2416.

Vargas, J., J. Oton, R. Marabini, S. Jonic, J. M. de la Rosa-Trevin, J. M. Carazo and C. O. Sorzano (2013). “FASTDEF: fast defocus and astigmatism estimation for high-throughput transmission electron microscopy.” J Struct Biol 181(2): 136–148.

Vargas, J., J. A. Quiroga, C. O. Sorzano, J. C. Estrada and J. M. Carazo (2011). “Two-step interferometry by a regularized optical flow algorithm.” Opt Lett 36(17): 3485–3487.

Vargas, J., J. A. Quiroga, C. O. Sorzano, J. C. Estrada and M. Servin (2012). “Multiplicative phase-shifting interferometry using optical flow.” Appl Opt 51(24): 5903–5908.

Vargas, J., R. Restrepo, J. C. Estrada, C. O. Sorzano, Y. Z. Du and J. M. Carazo (2012). “Shack-Hartmann centroid detection using the spiral phase transform.” Appl Opt 51(30): 7362–7367.

Vargas, J., R. Restrepo, J. A. Quiroga and T. Belenguer (2011). “High dynamic range imaging method for interferometry.” Optics Communications 284(18): 4141–4145.

Vilas, J. L., J. Gomez-Blanco, P. Conesa, R. Melero, J. Miguel de la Rosa-Trevin, J. Oton, J. Cuenca, R. Marabini, J. M. Carazo, J. Vargas and C. O. S. Sorzano (2018). “MonoRes: Automatic and Accurate Estimation of Local Resolution for Electron Microscopy Maps.” Structure 26(2): 337–344 e334.

Vilas, J. L., H. D. Tagare, J. Vargas, J. M. Carazo and C. O. S. Sorzano (2020). “Measuring local-directional resolution and local anisotropy in cryo-EM maps.” Nat Commun 11(1): 55.

Walls, A. C., Y. J. Park, M. A. Tortorici, A. Wall, A. T. McGuire and D. Veesler (2020). “Structure, Function, and Antigenicity of the SARS-CoV-2 Spike Glycoprotein.” Cell.

Wandzik, J. M., T. Kouba, M. Karuppasamy, A. Pflug, P. Drncova, J. Provaznik, N. Azevedo and S. Cusack (2020). “A Structure-Based Model for the Complete Transcription Cycle of Influenza Polymerase.” Cell.

Wang, Q., R. A. Corey, G. Hedger, P. Aryal, M. Grieben, C. Nasrallah, A. Baronina, A. C. W. Pike, J. Shi, E. P. Carpenter and M. S. P. Sansom (2020). “Lipid Interactions of a Ciliary Membrane TRP Channel: Simulation and Structural Studies of Polycystin-2.” Structure 28(2): 169–184 e165.

Winn, M. D., G. N. Murshudov and M. Z. Papiz (2003). “Macromolecular TLS refinement in REFMAC at moderate resolutions.” Methods Enzymol 374: 300–321.

Wrapp, D., N. Wang, K. S. Corbett, J. A. Goldsmith, C. L. Hsieh, O. Abiona, B. S. Graham and J. S. McLellan (2020). “Cryo-EM structure of the 2019-nCoV spike in the prefusion conformation.” Science 367(6483): 1260–1263.

Yang, M., Y. S. Chen, M. Ichikawa, D. Calles-Garcia, K. Basu, R. Fakih, K. H. Bui and K. Gehring (2019). “Cryo-electron microscopy structures of ArnA, a key enzyme for polymyxin resistance, revealed unexpected oligomerizations and domain movements.” J Struct Biol 208(1): 43–50.

